# A personalized, multi-omics approach identifies genes involved in cardiac hypertrophy and heart failure

**DOI:** 10.1101/120329

**Authors:** Marc Santolini, Milagros C. Romay, Clara L. Yukhtman, Christoph D. Rau, Shuxun Ren, Jeffrey J. Saucerman, Jessica J. Wang, James N. Weiss, Yibin Wang, Aldons J. Lusis, Alain Karma

## Abstract

Identifying genes underlying complex diseases remains a major challenge. Biomarkers are typically identified by comparing average levels of gene expression in populations of healthy and diseased individuals. However, genetic diversities may undermine the effort to uncover genes with significant but individual contribution to the spectrum of disease phenotypes within a population. Here we leverage the Hybrid Mouse Diversity Panel (HMDP), a model system of 100+ genetically diverse strains of mice exhibiting different complex disease traits, to develop a personalized differential gene expression analysis that is able to identify disease-associated genes missed by traditional population-wide methods. The population-level and personalized approaches are compared for isoproterenol(ISO)-induced cardiac hypertrophy and heart failure using pre- and post-ISO gene expression and phenotypic data. The personalized approach identifies 36 Fold-Change (FC) genes predictive of the severity of cardiac hypertrophy, and enriched in genes previously associated with cardiac diseases in human. Strikingly, these genes are either up- or down-regulated at the individual strain level, and are therefore missed when averaging at the population level. Using insights from the gene regulatory network and protein-protein interactome, we identify *Hes1* as a strong candidate FC gene. We validate its role by showing that even a mild knockdown of 20-40% of *Hes1* can induce a dramatic reduction of hypertrophy by 80-90% in rat neonatal cardiac cells. These findings emphasize the importance of a personalized approach to identify causal genes underlying complex diseases as well as to develop personalized therapies.

**Significance:** A traditional approach to investigate the genetic basis of complex diseases is to look for genes with a global change in expression between diseased and healthy individuals. Here, we investigate individual changes of gene expression by inducing heart failure in 100 strains of genetically distinct mice. We find that genes associated to the severity of the disease are either up- or down-regulated across individuals and are therefore missed by a traditional population level approach. However, they are enriched in human cardiac disease genes and form a coregulated module strongly interacting with a cardiac hypertrophic signaling network in the human interactome. Our analysis demonstrates that individualized approaches are crucial to reveal all genes involved in the development of complex diseases.

## Introduction

Contrary to “Mendelian” diseases where causality can be traced back to strong effects of a single gene, common diseases result from modest effects of many interacting genes (1). Understanding which genes are involved and how they affect diseases is a major challenge for designing appropriate therapies.

Heart failure (HF) is a well-studied example of a genetically complex disease involving multiple processes that eventually lead to a common phenotype of abnormal ventricular function and cardiac hypertrophy (2). Numerous studies have attempted to pinpoint differentially expressed genes (DEGs) to find biomarkers for the prognosis of the disease and the design of appropriate drugs (3), as well as explore underlying affected signaling pathways (4). Such studies typically compare gene expression between samples in healthy and diseased states, such as non-failing vs failing hearts in murine (5), canine (6), or human samples (see (7) for a broad review). However, because of the different genetic backgrounds of the surveyed individuals, as well as different severities of HF, those studies show very limited overlap of DEGs. While separate studies typically identify tens to hundreds of DEGs, not a single DEG is common to all studies (7). Moreover, it is unclear whether the healthy state is itself a well-defined unique state. In particular, several studies have shown that, due to compensatory mechanisms involved in homeostasis, different combinations of ion channel conductances in neurons and cardiac cells can lead to a normal electrophysiological phenotype, e.g. a similar bursting pattern of motor neurons or a similar cardiac action potential and calcium transient (8, 9). This has led to the concept that genetically distinct individuals represent different “Good Enough Solutions” corresponding to distinct gene expression patterns underlying a healthy phenotype. Different combinations of gene expression in a healthy state resulting from genetic variations would be expected to yield different DEGs in a diseased state. Thus, small numbers of DEGs that are only shared by a subset of individuals, and would be missed by a standard population-wide DEG analysis, could in principle have a causal role. Identifying these genes remains a central challenge in personalized medicine (10, 11).

In order to explore the variability of individual trajectories leading to hypertrophy and HF, we leverage the Hybrid Mouse Diversity Panel (HMDP), a model system consisting of >100 genetically diverse strains of mice that we described previously (12, 13). This model system provides an unprecedented opportunity to investigate the role of strain-to-strain genetic differences on phenotypic outcome by allowing us to measure strain-specific changes of gene expression in response to a disease-inducing stressor in a large number of strains under well-controlled environmental conditions. Moreover, those strain-specific findings can be replicated by studying the phenotype of several genetically identical mice from the same strain, thereby disentangling intra-strain and inter-strain variations. In the specific case of HF onto which we focus here, such data could not be obtained in human studies where heart tissue biopsies have been extracted from either donor hearts or explanted hearts of late stage HF patients in a genetically diverse population (14). Indeed, gene expression data obtained from those biopsies can only be used to perform a population-level differential gene expression analysis. In contrast, here we identify relevant genes by correlating strain-specific temporal changes of gene expression, i.e. gene expression measured genome-wide before and after a stressor, with the corresponding strain-specific changes of phenotype. The ability to study a large number of strains using the HMDP is essential to have enough statistical power to establish such a correlation, a power that has been lacking from previous studies limited to small numbers of strains (15-18).

We use both gene expression and phenotypic data acquired in HMDP strains before and three weeks after implantation of a pump delivering isoproterenol (ISO), a stressor inducing HF (12). In the HMDP, the pathological stressor induces a global response characterized mainly by cardiac hypertrophy along with more marginal changes in chamber dilation and contractile function at the population level. As a result, we primarily focus on the identification of genes relevant for cardiac hypertrophy. Expression data is collected at the whole heart level and the Total Heart Weight is used to quantify the degree of cardiac hypertrophy. Importantly, the severity of the hypertrophic response is highly variable among strains, ranging from almost no hypertrophy to up to an 80% increase of heart mass. Our study is directed at understanding why certain individuals are more susceptible to or protected against cardiac hypertrophy due to their genetic backgrounds.

In the following, we develop a personalized strategy to find genes for which the individual, strain-specific fold-change (FC) of expression is associated with the degree of hypertrophy. We find a small set of 36 genes that we refer to as FC genes. We then compare them to genes identified using Significance Analysis of Microarrays (SAM), a standard tool to evaluate population-wide DEGs (19). Interestingly, the FC genes are not identified as significantly changed at the population level. They indeed typically have opposite fold changes in low and high hypertrophy strains that cancel each other when averaged over all strains. We show that the FC genes are strongly enriched in cardiac disease genes from previous Genome-Wide Association Studies (GWAS), while SAM genes are in contrary enriched in fibrosis genes. We then show that those two sets form two distinct communities in the co-expression network among healthy as well as ISO-injected strains and we identify potential Transcription Factors (TFs) to explain the observed co-regulation of FC genes. Moreover, we find that the proteins encoded by the FC genes, but not the SAM genes, interact predominantly with proteins belonging to a cardiac hypertrophic signaling network (CHSN) that has been shown to provide a predictive model of hypertrophy in relation to multiple stressors including ISO (20). Interestingly, we find that one of the FC genes, namely *Hes1*, is also a predicted TF and an important interactor with the CHSN. Using a knockdown approach, we find that it plays a major role in cardiac hypertrophy, allowing us to validate our personalized, multi-omics approach.

## Results

### 1. Two types of responses to stressor-induced cardiac hypertrophy and heart failure

We begin with an example showing two distinct ways to describe the response to ISO in the HMDP (Figure 1). First, one can note that ISO induces a global response across all strains, resulting in cardiac hypertrophy. This is seen in Figure 1a, where the distribution of heart mass among the post-ISO strains can clearly be distinguished from the pre-ISO distribution (p < 2.2e-16 under Student t-test). At the gene level, such a response is typically analyzed by looking for DEGs at the population level, i.e. genes for which the change in average expression with the stressor is significantly greater than the variability with and without the stressor (Figure 1b). Typical tools include t-test (21), SAM (19), or LIMMA (22). Genes found with these methods have a differential expression profile at the population level and are therefore potential biomarkers of the trait of interest (see microarray data for *Serpina3n* in Figure 1c). However, despite the global response in the level of gene expression to ISO, the degree of hypertrophy among individual strains is highly variable, from almost none to an 80% increase of heart weight (Figure 1d). This calls for an evaluation of the strength of differential gene expression at the individual level. In particular, a whole new class of genes becomes available for analysis. Indeed, even if a gene does not show population-wide average differential expression, it can show extensive variation at the individual, strain-specific level (Figure 1e). This is the case for the gene *Kcnip2* encoding the protein KChIP2, which interacts with pore forming subunits (Kv4.2 and Kv4.3) of the transient outward current I_to_ expressed in heart, and which has been implicated in cardiac hypertrophy (23-25). Though not showing population-wide differential expression (Figure 1f), its individual fold-change of expression can vary drastically from 2-fold decrease to a 2-fold increase depending on the considered strain (Figure 1g). Interestingly, when comparing the individual variations of those two types of genes with the degree of hypertrophy (Figure 1h,i,k), one can see that global DEGs are not necessarily good descriptors of the individual changes of phenotype (Figure 1j), unlike the second type of genes missed by a traditional population-wide method (Figure 1l). In particular, in the case of *Kcnip2*, we observe a significant positive correlation with the severity of hypertrophy (r=0.4, p=1.5e-4). This is particularly interesting since *Kcnip2* has previously been shown to be down-regulated during cardiac hypertrophy (24, 26) in the strain 129X1/SvJ. While we confirm this finding, we also observe that it is unusual in a broader context, and that *Kcnip2* is most of the time up-regulated in strains with marked hypertrophy.

**Figure 1.**
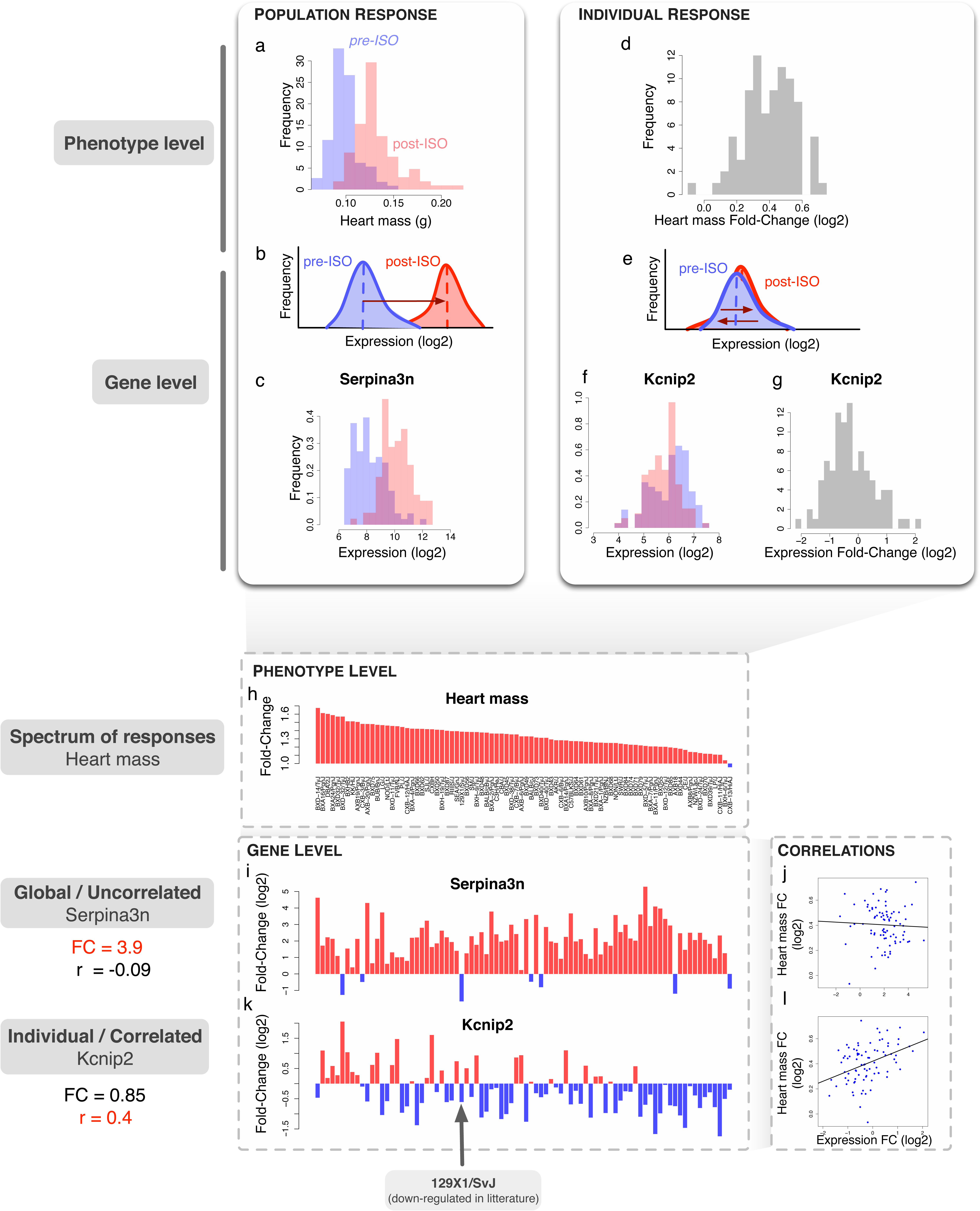
Two types of responses to stressor-induced heart failure. a. Histograms of the pre-ISO (blue) and post-ISO (red) heart masses of the HMDP strains.
b. Typical Differentially Expressed Genes (DEGs) show clear population-average fold-change allowing distinguishing the two populations of strains.
c. An example of such strong DEG, namely *Serpina*3n.
d. Histogram of the heart mass fold-change (FC) computed for each strain from the HMDP.
e. Expression FC at the individual level can lead to cases were the population-average FC is null while the individual FCs are not.
f. *Kcnip2* is a good example of a gene with no population-wide average FC.
g. However, at the individual level, *Kcnip2* shows strong variations, as seen in the histogram of individual FCs at the strain level (log2 of post over pre-ISO expression ratio). In particular, some strains have a 4 fold decrease of expression (-2 in log2), while others have a 4 fold increase (+2).
h. For better visualization, the strain-specific heart mass FC is shown by decreasing strength. Red bars indicate increase and blue bars decrease in value.
i. *Serpina*3n log FC is shown with the same strain ordering than in h. Its population-wide FC is high (3.9), with most strains showing a strong positive FC (red bars).
j. However, the correlation of *Serpina*3n FC with the heart mass FC is not significant (r=-0.09, p=0.43).
k. On the other hand, *Kcnip2* shows a weak population-wide FC (FC=0.85). In particular, some strains show an increased expression (red bars) while others show a decreased expression (blue bars). The red arrow indicates the 129X1/SvJ strain in which *Kcnip2* has previously been shown to be down-regulated during cardiac hypertrophy (24).
l. Contrary to *Serpina*3n, *Kcnip2* FC is significantly correlated to heart mass FC (r=0.4, p=1.5e-4), with increased expression corresponding to high hypertrophy and decreased expression corresponding to low hypertrophy.

In the following, we generalize these observations to identify a larger set of genes that, like *Kcnip2*, have an individual FC correlated to the severity of hypertrophy, and we compare this set to the complete set of DEGs identified by the population-level SAM method.

### 2. Identification of genes associated to the severity of hypertrophy

Here we develop a method to determine which genes show individual, strain-specific expression FCs significantly correlated to the individual hypertrophic response measured by the individual fold-change of heart mass. We use microarray and phenotype expression data described in (13). Since our methodology is based on correlations, we choose to select those genes that belong to the giant component of the gene co-expression network above a certain correlation cutoff (see Methods and Figures S1,2). The advantages of such a filter compared to one based on absolute expression levels is that it yields a clear, well-defined cutoff (Figure S1b) while also rejecting genes having high expression but artefactual correlations (e.g. hitting the microarray saturation level in Figure S1c). We obtain a filtered set of 11,279 high-confidence genes. We then compute for all genes the absolute Pearson correlation between the gene expression fold-change and the individual hypertrophic response (Figure 2a, blue histogram). To control for False Positives, we compute the expected correlations when randomizing the phenotype by shuffling strain labels (see Methods and Figure 2a, red histogram). One can see significant enrichment in genes with high correlation to the trait. To quantify this enrichment, we compute the proportion of observed (‘blue’) correlations divided by the proportion of correlations in the randomized cases (‘red’) above various correlation cutoffs. Figure 2b shows this enrichment as a function of the gene rank, ordered by decreasing absolute value of the correlation with hypertrophy. The enrichment shows a peak at 36 genes, followed by a plateau until ∼500 genes, and a subsequent decrease. We define these 36 genes as our candidates to describe the hypertrophic spectrum. These genes are listed in Table 1, along with references supporting the involvement of several of them in cardiac hypertrophy and HF. In the following, we refer to this set of genes as the “FC” set. The full list of genes with their correlation with hypertrophy is provided in Supplementary Table S1.

**Figure 2.**
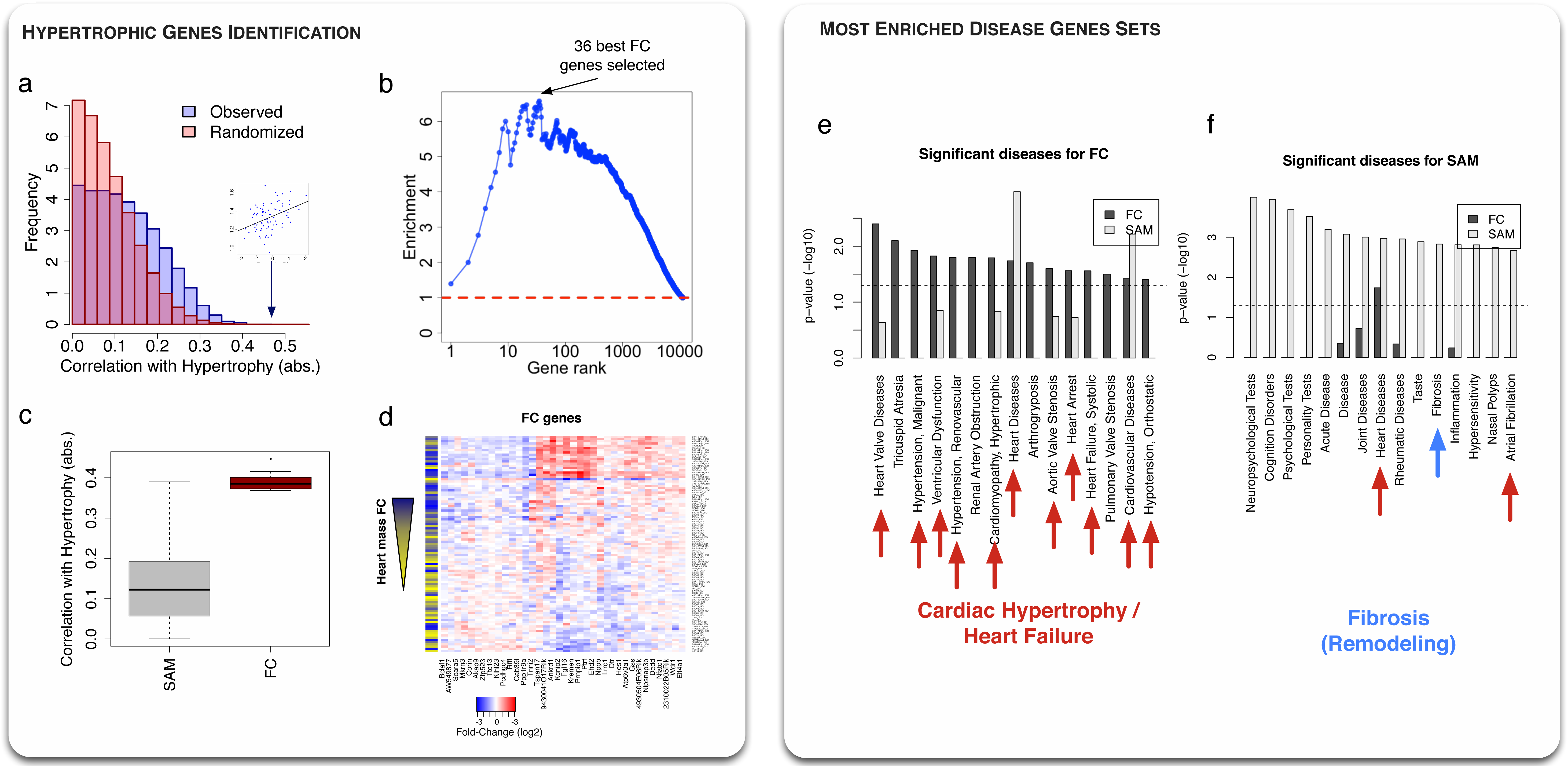
Identification of genes associated to the severity of cardiac hypertrophy. a. Histogram of the absolute values of the correlations between the FCs of genes expression and hypertrophy for all genes (blue, observed, red, randomized phenotype). Genes individual FCs are more correlated to hypertrophy than expected. Inset plot corresponds to the best observed correlation.
b. The previous enrichment is assessed by computing the ratio of the area under the observed and randomized curve as a function of correlation cutoffs. Cutoffs are matched to the genes correlations ranked in decreasing order. The enrichment peaks at N=36 genes, which defines the set of “FC” genes.
c. Boxplot comparing the values of the absolute correlation with hypertrophy for the 2,538 SAM genes resulting from a population-wide DEG study (see main text) and for the 36 identified individual FC genes. FC genes have significantly higher correlation.
d. Heatmap showing the 36 genes (columns) log fold-changes across strains (rows). The left column shows the degree of hypertrophy (yellow=low, dark blue=high). Hierarchical clustering shows a natural grouping of the strains by the severity of hypertrophy.
e. Enrichment of 36 best FC genes in human disease genes from GWAS studies. The 15 most enriched sets are shown. Red arrows indicate cardiac diseases (11/15). The enrichment of the 36 best SAM genes is shown for comparison, with low enrichment in the found sets.
f. Similar than g, for 36 SAM genes. These genes show enrichment in “Fibrosis”, a feature of structural remodeling during cardiac hypertrophy.

**Table 1.**
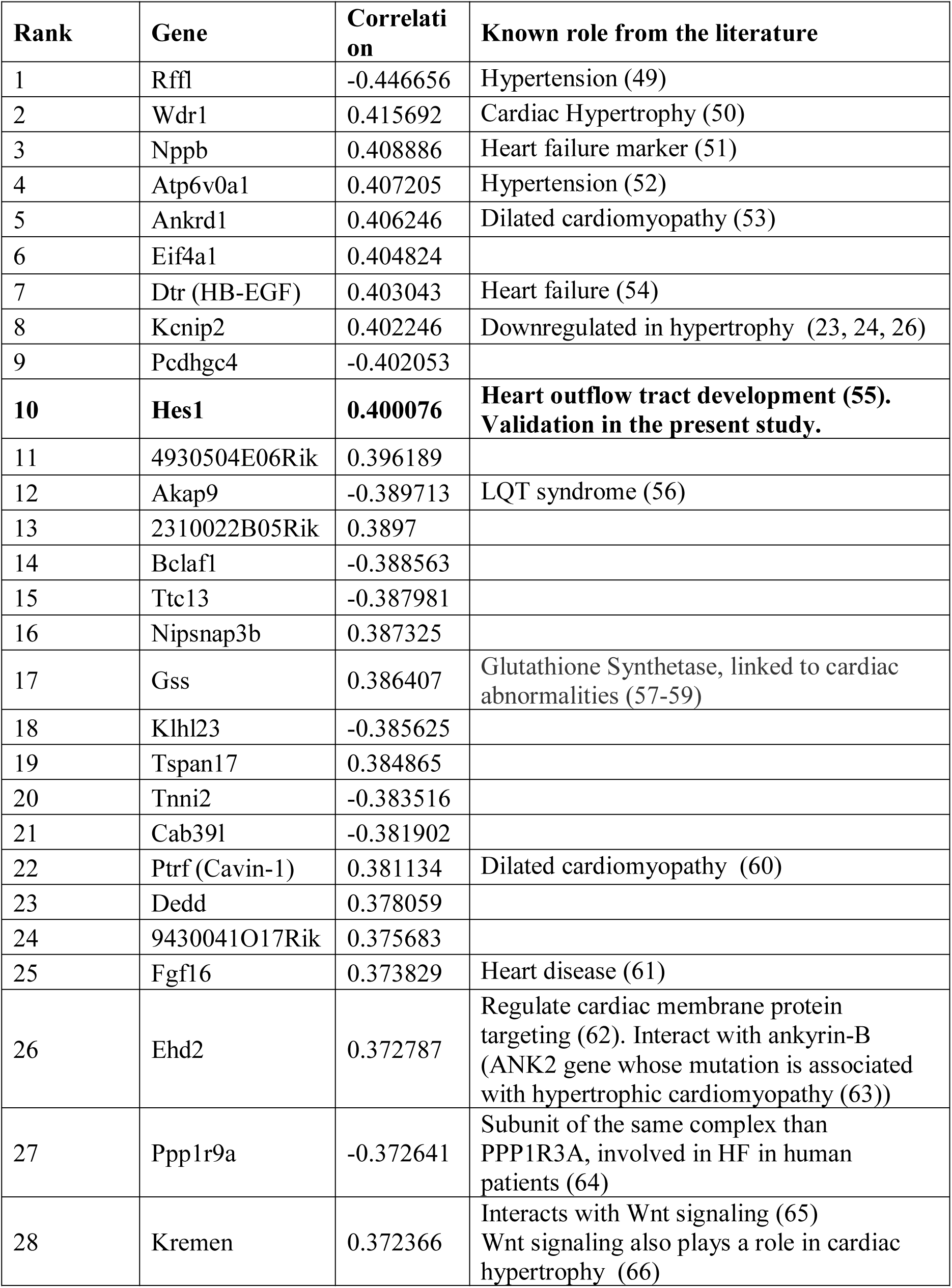
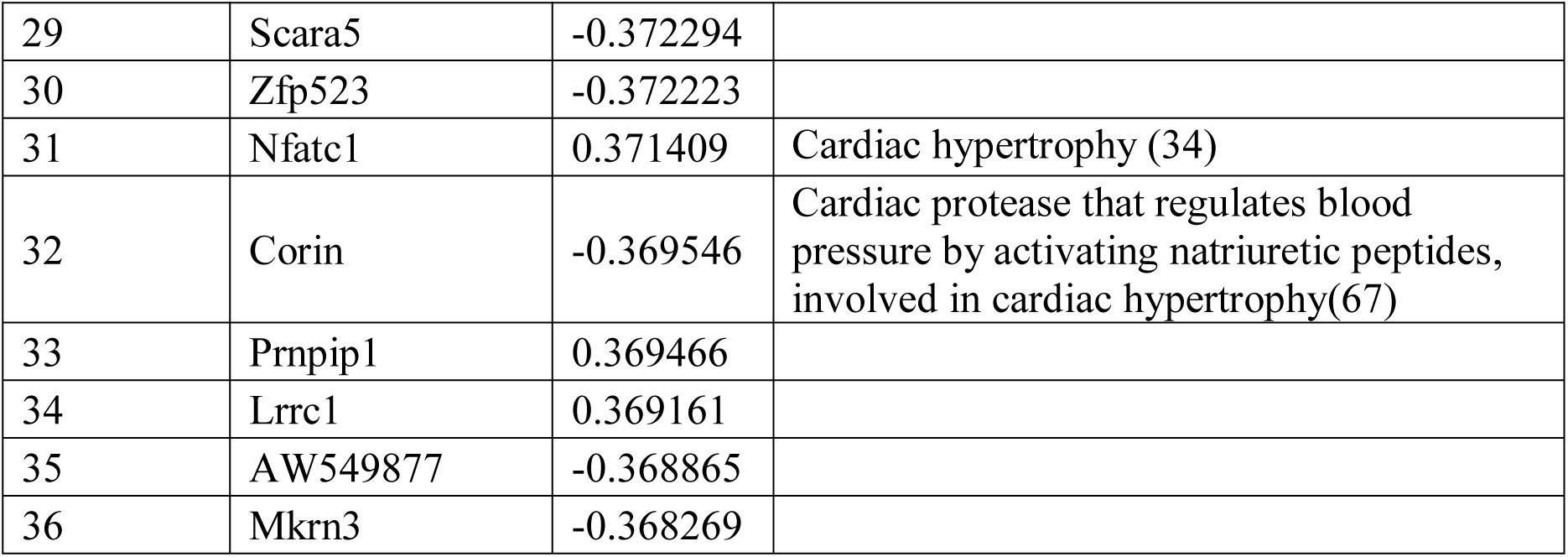
List of genes predicted with the individual Fold-Change analysis.

As a comparison, we compute the population-wide DEGs using Significance Analysis of Microarray or SAM (19). This exhibits 2,538 DEGs at a False Discovery Rate of 1e-3 (see Methods). Interestingly, we find no significant overlap (p=0.68, hypergeometric test) between these SAM genes and the FC set, with 6 genes common to both sets (*Tspan17, Ppp1r9a, Bclaf1, AW549877, Gss*, 2310022B05Rik and 9430041O17Rikm). In general, correlations between the individual fold-changes of the SAM genes and the degree of hypertrophy are found to be quite low (Figure 2c).

The 36 FC genes are shown in Figure 2d. As expected from the absence of overlap with SAM genes, the FC genes have both negative (blue) and positive (red) fold-change across the different strains, meaning that they have negligible average fold-change at the population level. A question that arises is whether the variability observed in the individual fold-changes of gene expression across strains is a consequence of genetic variability, or merely reflects environmental or experimental spurious effects. To investigate this question, we take advantage of the fact that expression data has been replicated in 8 strains, either in pre-ISO, post-ISO or both. Since mice from the same strain have a similar genetic background, they should therefore show very comparable individual fold-changes. Expression fold-change is shown for the 36 FC genes for the replicated strains in Figure S3a. We assess the replicability by computing the Spearman rank correlation of the 36 FC genes fold-change profiles between mice from replicated strains. We find a mean correlation of 0.76, compared to 0.14 for pairs of strains taken at random among the non-replicated pool (p = 1.6e-7, Wilcoxon test, see Figure S3b). This result shows that individual fold-changes are tightly controlled at the genetic level.

In the following, we wish to evaluate further the biological signal carried by these FC genes missed by population-wide methods.

### 3. Biological relevance of the identified FC genes

Given the importance of the genetic control of those genes, they must be more susceptible to genetic variations. To explore that idea, we look at the enrichment in disease genes coming from previous GWAS. We use HuGE database of human genes associated to 2,711 different diseases (see Methods). First, we convert the mouse gene names to human as described in the Methods. Then, we rank the diseases according to their enrichment in 36 FC (resp. 36 best SAM) genes using a hypergeometric test assuming as null hypothesis a uniform repartition of the genes across diseases. Results are shown in Figure2e,f for the 15 most enriched diseases in each case. We observe that FC genes are strongly enriched in heart diseases (11 in the 15 most enriched diseases) while SAM genes are only enriched in two cardiac diseases and in fibrosis, a feature characteristic of the structural remodeling taking place during HF (27). Those findings exhibit two distinct roles of FC and SAM genes in the progression of cardiac hypertrophy. While the cross-talk between cardiac fibroblasts and myocytes during cardiac hypertrophy has been studied previously (28), here we disentangle their relative contributions into a shared, population-wide fibroblastic component, and a fine-tuned, individualized component capable of explaining the severity of cardiac hypertrophy. Moreover, the enrichment of FC genes in human GWAS genes also highlights the relevance of the present HMDP data analysis to human cardiac hypertrophy and HF.

### 4. Co-expression and co-regulation

The identified sets of population-level and individual FC genes have until now been considered as collections of independent genes. However, in the cell, genes function together to achieve higher-order physiological functions. Such a collective behavior can be assessed in the framework of co-expression networks, where genes are related by the similarity of their profile of expression across different conditions. In the context of the HMDP, we investigate whether the predicted sets of genes show evidence of co-regulation in healthy and post-ISO hypertrophic strains. To that extent, we compute the squared Pearson correlations (r^2^) between the 36 best genes of both the FC and SAM sets. Correlation matrices are then cut off at r^2^>0.1 to keep significant interactions. We show in Figure 3a and b the resulting co-expression networks in pre and post-ISO conditions. We clearly see that the two sets of genes form dense modules, and are disconnected from each other, with only few links between the two sets. Interestingly, we see that the biomarker and modulator of hypertrophy *Nppb* (29) acts as a bridge between the two modules in pre-ISO condition (Figure 3a, top), and is even found strongly co-expressed with the SAM genes in post-ISO mice (Figure 3b). This suggests a role for *Nppb* in driving a cross-talk between FC genes and SAM genes. Finally, to quantify the relative density of the modules, we compared them to 1,000 sets of a similar number of randomly selected genes. We show the resulting Z scores in Figure 3c. Both SAM and FC sets show much stronger co-expression than randomly expected, with the SAM module being even denser under ISO condition. On the contrary, the density of links between the two modules is significantly smaller than expected by chance, indicating that the two sets of genes are disjoint sets in the co-expression network. Overall, these results show that the FC and SAM genes form two tight, disjoint communities in the co-expression network, both in pre-ISO and post-ISO mice.

**Figure 3.**
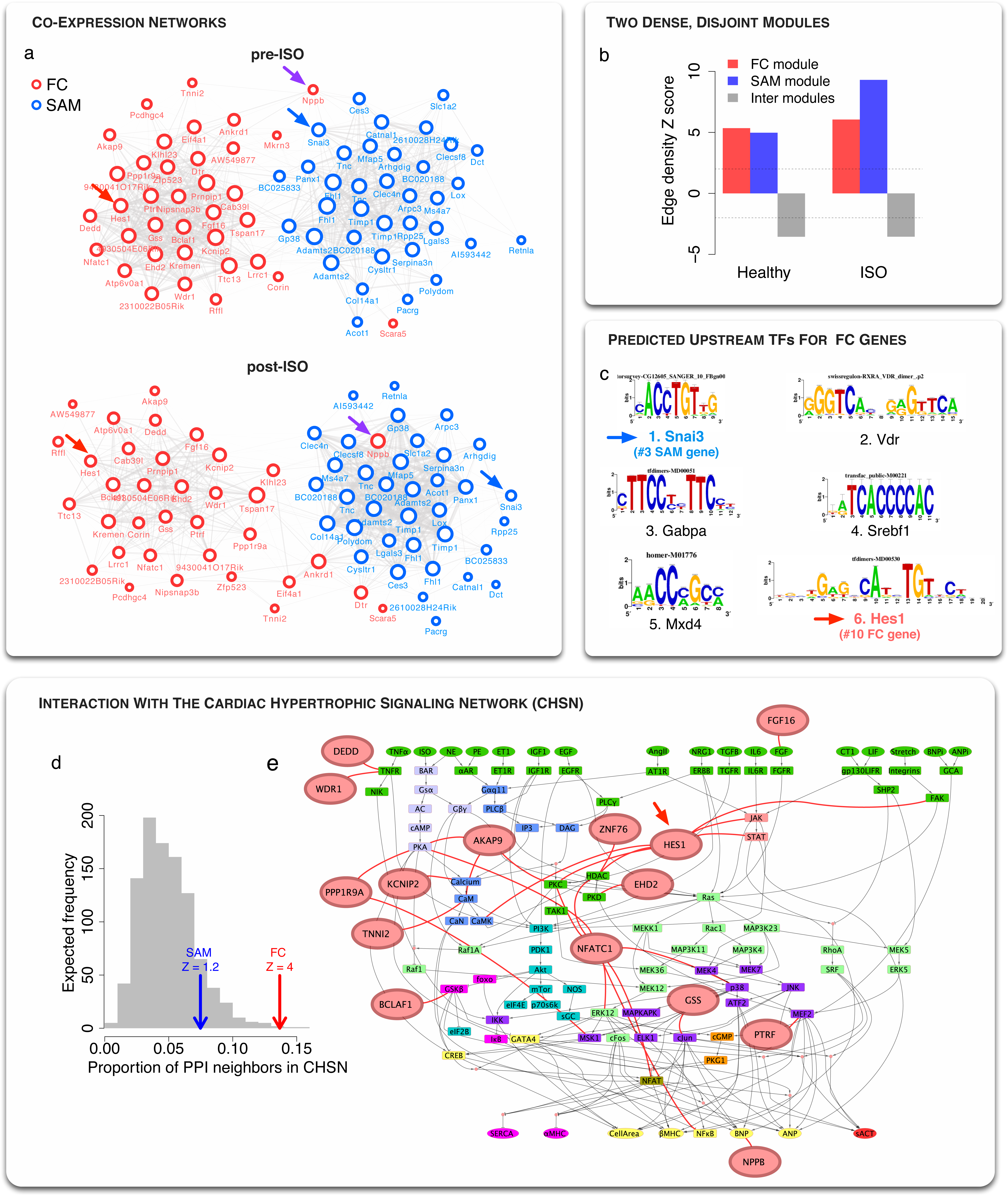
FC genes are co-regulated and significantly connected to the cardiac hypertrophy signaling network (CHSN) a. Co-expression networks of the 36 best FC and SAM genes in healthy and post-ISO hypertrophic strains. Edges are drawn between two genes if the square Pearson correlation is greater than 0.1 (r^2^>0.1). The two modules segregate naturally using a force layout algorithm, showing that the modules have high clustering but only few links between themselves. Interestingly, Nppb (purple arrow) segregates with SAM genes, especially in ISO condition.
b. The edge density of the FC module, the SAM module, and the FC to SAM edges is computed and compared to the density expected for random sets of nodes of the same size (see Methods). The corresponding Z scores are significantly high (Z > 2) for both modules, indicating high co-expression. However, there are significantly fewer links than expected between the two modules (Z < -2), indicating that they are disjoint in the coexpression network,
c. List of the 6 most enriched TF motifs in the -/+20kb regions around the 36 FC genes TSSs predicted using iRegulon (30). Interestingly, Snai3 (blue arrow) is a SAM gene and Hes1 (red arrow) a FC gene, suggesting a crosstalk between the two modules at the gene regulatory level.
d. Proportion of neighbors in the interactome that belong to the Cardiac Hypertrophy Signaling Network or CHSN (20) for different gene sets: the FC set (red arrow), the 36 best SAM genes (blue arrow) and 1,000 realizations of random nodes in the interactome with the same size as the FC set (gray histogram). Z scores are computed relative to the gray distribution. The FC set is significantly connected to the CHSN, while the SAM genes are not significantly different than a random set.
e. Network visualization of the CHSN (48), along with neighbors from the 36 best FC genes (red nodes). A more detailed interaction network is shown in Figure S5.

The finding that the FC genes are strongly co-expressed suggests that they are co-regulated. To explore this possibility, we look for enrichment in common TF binding sites in the vicinity of the 36 FC genes. To compute the enrichment, we use iRegulon, a recent algorithm integrating different TF motifs databases and using phylogenic conservation to identify overrepresented binding sites in the -20/+20kb regions around the Transcription Start Sites of genes of interest (see Methods) (30). The identified motifs are then ranked by target enrichment among selected genes, and are associated to a list of putative TFs that can bind them (Figure 3c). The full list of predictions is given as Supplementary Table S2. We find that the best-ranked motif is associated with repressor TFs Scrt1 and Scrt2, known to modulate the action of basic helix-loop-helix (bHLH) TFs (31). Interestingly, the corresponding PWM motif is also matched to Snai3 TF, a gene ranked 3^rd^ among SAM genes. The 2^nd^ motif, VDR, is known to be involved in heart failure and cardiac hypertrophy (32). Finally, the sixth predicted TF is associated to Hes1, which ranks 10^th^ among the FC genes. This indicates that there is a cross-talk between the two modules at the gene regulatory level, with both FC and SAM genes being involved in the regulation of the expression of the FC genes.

### 5. Exploration of the neighborhood in the interactome

While useful to detect gene regulatory changes involved in the disease process, gene expression does not capture post-translational changes and interactions that occur at the protein level. To explore the potential involvement of the predicted sets of genes at the protein level, we use a previously published human interactome combining high-throughput and literature curated protein-protein, metabolic, kinase-substrate, signaling and to a lesser extent regulatory interactions (33). After converting to human gene symbols (see Methods), the proteins encoded by the 36 best FC and SAM genes have respectively 364 and 346 interacting partners. We then compute pathway enrichment for these neighbors (see Methods). The other most highly enriched pathway is linked to NFAT signaling, known to be important in HF (34). Interestingly, we find that the second most enriched pathway for FC neighbors is a previously published Cardiac Hypertrophy Signaling Network (CHSN) containing 106 nodes (corresponding to 218 genes) giving a *predictive* model of hypertrophy in response to multiple stressors including ISO (20) (Figure S4). Indeed, about 14% of FC neighbors are components of this network, compared to a predicted random association of 4% (Z=4, Figure 3d). The CHSN is shown in Figure 3e and in more details in Figure S5, along with FC nodes directly interacting with CHSN nodes. In particular, we find that Hes1 is interacting with several nodes of the CHSN at different levels of the hierarchy, namely FAK, JAK, STAT, CamK, PKC and HDAC.

### 6. Experimental validation of Hes1

The previous results point toward a role for *Hes1* in cardiac hypertrophy and heart failure. Indeed, *Hes1* was found to be a FC gene, an upstream regulator of FC genes, and an interactor with several components of the CHSN. To determine the function of *Hes1* in the context of cardiac hypertrophy and heart failure, we performed siRNA knockdown in neonatal rat ventricular myocytes (NRVMs) followed by treatment with beta-adrenergic agonist isoproterenol (ISO) or alpha-adrenergic agonist phenylephrine (PE) containing media. Both agents induce hypertrophy through different molecular pathways, as can be seen in the CHSN (see Figure 3e). Using siRNA to silence *Hes1* expression, we achieved a 20% to 40% decrease in *Hes1* expression when compared to transfection control (Figure 4a). At the molecular level, treatment with either ISO or PE containing media drastically increases the expression of the HF markers *Nppa* and *Nppb*, which rose 3.5- and 7.9-fold, respectively under ISO treatment and 11-fold and 13-fold, under PE treatment in cells transfected with the control siRNA. Strikingly, knockdown of *Hes1* expression strongly impaired the induction of these two markers under both treatment conditions. *Nppa* induction was reduced up to 110% and 88% under ISO and PE treatment while *Nppb* induction was reduced up to 66% and 91% under ISO and PE treatment, respectively. In addition to these molecular changes, we investigated the role of Hes1 in modulating the increase in cardiomyocyte cell size upon treatment with ISO and/or PE. As expected, following ISO/PE treatment, cells transfected with the control siRNA doubled in cellular cross-sectional area (Figure 4c and Figure S6). In comparison, cells transfected with the *Hes1* siRNA showed up to 87% and 79% reduction in cell size increase following treatment with ISO and PE, respectively. This effect is consistent with the fact that HMDP strains showing no or mild hypertrophy exhibit strong negative fold-change of *Hes1* (Figure S7). Taken together, these findings strongly suggest a role for *Hes1* as a novel regulator of cardiac hypertrophy *in vitro*.

**Figure 4.**
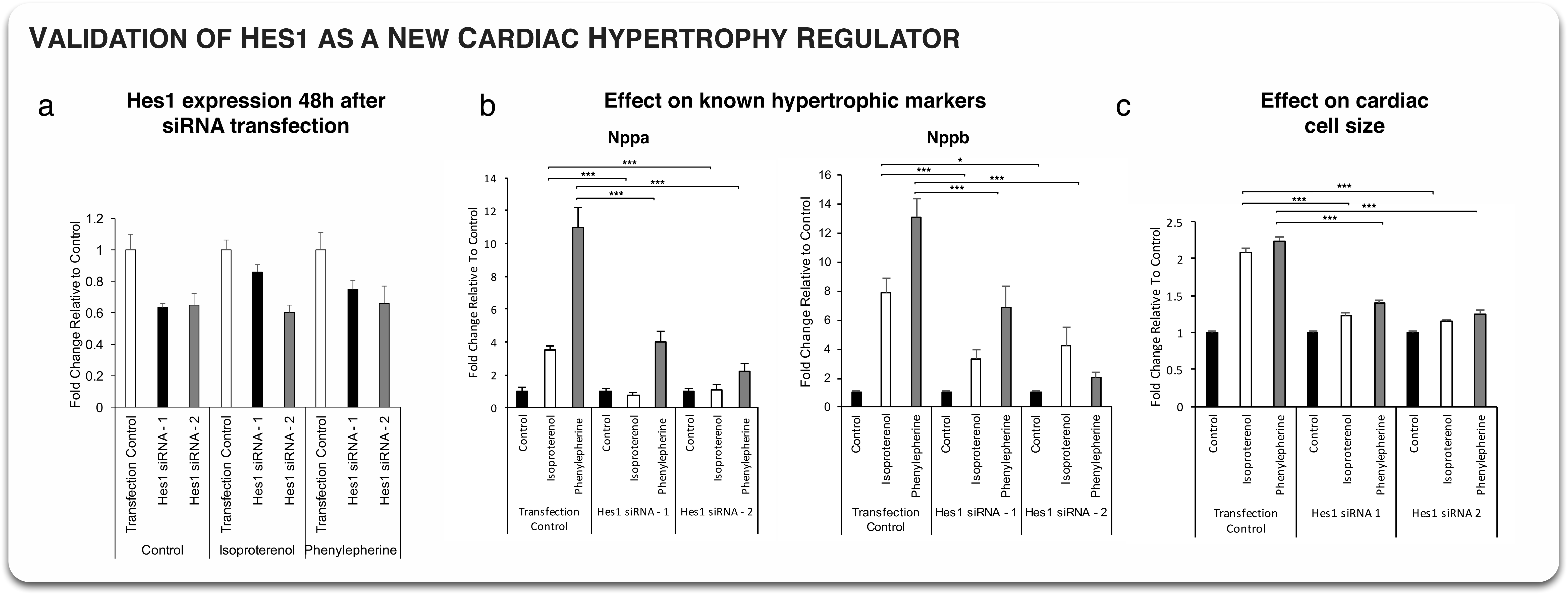
Validation of Hes1 as a new cardiac hypertrophy regulator. a. *Hes1* mRNA expression following 48h after siRNA transfection in a control, isoproterenol or phenylephrine medium. Three siRNAs were used, a scrambled, control one and two *Hes1* specific siRNAs. Both *Hes1* siRNAs show systematic downregulation of *Hes1* mRNA in all conditions.
b. Effect of *Hes1* knockdown on the known hypertrophic makers *Nppa* and *Nppb*. In both case, *Hes1* knockdown leads to a significant change in biomarkers activation in isoproterenol and phenylephrine conditions (* p<0.05, *** p<1e-3, Student t-test).
c. Effect of Hes1 knockdown on cardiac cell size relative to control medium cell size. Both siRNAs lead to a drastic 80-90% decrease in hypertrophy in both isoproterenol and phenylephrine media.

### 7. Discussion

In the present study, we investigated the spectrum of cardiac hypertrophy and HF development in 100+ genetically diverse mice from the HMDP when subjected to chronic ISO infusion. We have analyzed two types of responses. First, the global response at the population level with a large number (1,000+) of genes involved, as detected by the SAM algorithm. Their global fold-change is representative of the global hypertrophy observed across all strains. However, the magnitude of their fold-change at the individual level does not predict the degree of individual hypertrophy. Using a correlation-based method, we found another group of ∼40 genes that predicts the degree of hypertrophy. We named these the “FC” genes in reference of the fact that we found them using their individual, strain-specific fold-change. Surprisingly, these genes have a near zero fold-change at the population level due to the cancelling contributions of up- and down-regulation in different strains, so that they are not detected using classical differential expression tools. While several FC genes have previously been implicated in cardiac hypertrophy and HF (see Table 1), their high variability in such a controlled setup has not been explored previously. We showed that these genes are enriched for heart failure gene candidates previously described in the literature, as well as for human cardiac disease genes. On the other hand, the best SAM genes are enriched in fibrosis disease genes. ISO has been shown to induce first myocardial fibrosis concomitantly with myocyte necrosis, followed by myocyte hypertrophy on a longer time scale (35), and fibrosis is also known to be an early manifestation of hypertrophic cardiomyopathy (36). Our results suggest that population-level SAM genes are predominantly associated with the early fibroblast response. On the other hand, since the change of heart mass is primarily determined by myocyte growth, our results suggest that FC genes are associated with the strain-specific degree of myocyte growth induced by beta-adrenergic stimulation.

We further investigated the roles of these genes in different biological networks. We found that both FC and SAM genes form distinct co-expressed modules. Interestingly, *Nppb* (encoding the BNP protein), a widely used biomarker and modulator (29) of HF, belongs to the FC set but is co-expressed with SAM genes in healthy mice, providing a unique bridge between the two sets. We note that this result is consistent with the previous finding that *Nppb* is an antifibrotic hormone produced by myocytes with an important role as a local regulator of ventricular remodeling in mice (37). Indeed, *Nppb* is correlated to the fibrotic SAM genes in healthy mice, consistent with a regulatory homeostatic behavior, but is found among FC genes after beta-adrenergic stimulation, consistent with a response proportionate to myocytes hypertrophy. It is also interesting to note that the SAM module overlaps significantly (p=3.4e-6, hypergeometric test) with a co-expression module previously found in post-ISO mice and shown to be involved in cardiac hypertrophy (38). Indeed, it shares the genes *Timp1, Tnc, Mfap5, Col14a1* and *Adamts2*, the latter of which was validated experimentally as a regulator of cardiac hypertrophy.

We then predicted several TFs to study this co-regulation. Interestingly, among the top TFs predicted as regulators of the FC genes, one of them, *Hes1*, belongs to the FC genes, and another one, *Snai3*, belongs to the SAM genes. We note that both inhibitory (Snai3, Hes1) and activatory (Vdr, Srebf1) TFs were found to have enriched binding sites around FC genes TSSs. This suggests a potential regulatory balance that could explain the up- and down-regulation observed for these genes across strains. We then looked at potential post-translational effects at the protein level by using an integrated interactome. We found that FC genes were strongly interacting with a cardiac hypertrophy signaling network (CHSN) previously shown to be predictive of cardiac hypertrophy in response to ISO and other stressors (20). This may indicate that several of those genes are upstream of a causal chain of events at the post-translational level that control myocyte growth. We note that the FC gene *Nppb* is present both as an input and an output of the CHSN. This exemplifies an interesting feedback architecture where downstream effects can causally affect upstream regulation. Overall, the FC genes constitute a HF “disease module” formed of co-regulated genes connected to the CHSN at the protein level.

A key finding of our study is that there is strong strain-to-strain variation in response to a stressor under similar well-controlled environmental conditions. This variation is largely explained by the different genetic backgrounds, as shown by the consistent responses in mice from same strains (Figure S3) and the strong enrichment in heart diseases GWAS (Figure 2e). For example, *Kcnip2* is known to be downregulated concomitantly with a reduction of Ito magnitude in cardiac hypertrophy (24, 26). Our results are consistent with this finding for the previously studied 129X1/SvJ strain (24), but show that *Kcnip2* is upregulated in many strains with pronounced hypertrophy leading to an overall positive correlation between *Kcnip2* expression and heart mass FC. This indicates that there are multiple possible compensatory mechanisms underlying a similar patho-phenotype. Similarly, we observed strong variation in the fold-change of *Nppb*. It was previously shown to be over-expressed during cardiac hypertrophy as an anti-fibrotic factor (29). Using our multiple strains setup, we observed a positive correlation between *Nppb* change of expression and the degree of hypertrophy. However, we also observed some cases were hypertrophic strains exhibit down-regulation of *Nppb*, including the widely used C57BL/6J and 129X1/SvJ strains (see Figures 2d and S3).

Finally, our approach was validated by testing *Hes1’s* role in cardiac hypertrophy. *Hes1* was chosen because of its involvement at different levels: found as a FC gene, *Hes1* is also a predicted TF regulating the FC genes and a key interactor of the CHSN. *Hes1* is part of the Notch signaling pathway which is highly conserved and involved in cell-cell communication between adjacent cells (39). This pathway is well known to play a crucial role in cardiac development and disease. Notch activity is required in complex organs like the heart that necessitate the coordinated development of multiple parts (40). Specifically, functional studies have shown that Notch activity is required for cardiovascular development and that Notch signaling causes downstream effects such as cell fate specification, cell proliferation, progenitor cell maintenance, apoptosis, and boundary formation (39). In previous studies, *Hes1* expression was observed to increase following myocardial infarction and other ischemic cardiomyopathies. Increased expression of *Hes1* was also shown to inhibit apoptosis of cardiomyocytes and promote instead their viability. However, whether *Hes1* acts as a regulator of novel heart failure markers has remained unclear (41). Here, we show that even a mild knockdown of 20-40% of *Hes1* can induce a dramatic reduction of hypertrophy by 80-90% (Figure 4c), identifying for the first time *Hes1* as a key regulator of cardiac hypertrophy. Importantly, this result is consistent with the HMDP data, where strains with no or mild hypertrophy have 20-50% decrease in *Hes1* after ISO injection (Figure S7b).

Overall, we have explored the individual, strain-specific responses to stressor-induced HF and identify 36 FC genes that are missed by traditional population-wide method of DEG analysis. We have shown that these FC genes provide a completely distinct, albeit complementary, picture of HF than population-wide DEGs. In particular, FC genes are enriched in human cardiac disease genes and hypertrophic pathways. This is important since previous studies that use population-level methods to identify DEGs have concluded that murine models are of limited relevance to human HF (42, 43). In contrast, our findings show that FC genes, identified by a personalized differential expression analysis in a genetically diverse population of mice, are relevant to human HF. By linking those genes both to upstream regulators and to a signaling network predictive of cardiac hypertrophy, we provide new insights into the regulation of the severity of and resistance to cardiac hypertrophy at the individual level, and validate *Hes1* as a novel regulator of cardiac hypertrophy *in vitro*. We believe this approach to be critically important for the appropriate design of upcoming experiments directed at unraveling causal genes in complex diseases.

## Methods

### Obtention of the data

Microarray data may be accessed at the Gene Expression Omnibus using accession ID: GSE48760. All phenotypic and expression data may also be accessed at https://systems.genetics.ucla.edu/data/hmdp_hypertrophy_heart_failure.

### Pre-filtering of the data

In order to reduce false positive predictions and computational time, we first filtered the 25,697 genes expression data. Instead of setting an arbitrary cutoff based on the level of expression as is commonly done, we decided to use a network approach that is consistent with the correlation-based methods used in this study. The idea is that the different genotypic backgrounds across strains lead to global gene expression modulation, thus creating correlation between expressed genes. Genes not associated to the core of varying genes should be the ones that carry too much experimental noise due to low expression or systematic biases.

We first computed the absolute Pearson correlation of gene expression fold-change between all pairs of genes. This creates a complete weighted network containing all genes. We then reasoned that genes for which expression is noisy because of low expression or experimental artifacts should have a low association to the other genes. We therefore looked at the size of the Largest Connected Component (LCC) of the network when hard-thresholding with several correlation cutoffs (figure S1a). We observed a fast decrease of the LCC size at low thresholds of 0.35-0.45, followed by milder steady decrease. The derivative of this curve is presented in figure S1b, showing a strong initial trough corresponding to noisy “satellite” nodes being cut from the LCC, followed by stabilization. We chose a cutoff of 0.5 corresponding to that stabilization plateau and kept the 11,279 genes in the LCC. The effect of this filter is made clear by looking at a selection of functional genes linked to the electromechanical coupling in heart cells (figure S1c). The rejected genes (gray bars) have either low expression (eg *Calm4, Kcnd3*) or display systematic saturation effects inherent to the microarray assay, which results in noisy correlations (eg *Tnnc1, Atp2a2*). More generally, we show in Fig S2 that filtered out genes show a correlation profile with hypertrophy similar to the one expected at random. In this paper, we use these 11,279 genes as input to the different methods.

### Computation of randomized correlations

To compute the expected correlations of Figure 2a, we first shuffle the heart mass fold-changes among strains. We then compute the correlations between all genes FCs and this randomized phenotype. We repeat that step 1,000 times. The final histogram is the average over the 1,000 randomizations.

### Computation of population-wide DEGs

The population-wide DEGs are computed by using Significance Analysis of Microarray or SAM (19) between the post-ISO and the pre-ISO expression data. Using a False Discovery Rate of 1e-3, we find 2,538 significant DEGs.

### Conversion from mouse symbols to human entrez IDs

In order to compute pathway and disease genes enrichment, we first needed to compute a table converting mouse gene symbols to human entrez IDs. We used UCSC genome browser mm9.kgXref, mm9.hgBlastTab and hg19.kgXref conversion tables available on the mySQL host genome-mysql.cse.ucsc.edu. The kgXref tables were used for conversion between symbols and entrez IDs while the Blast table was used to get the human orthologs of mouse genes.

### HuGE database

Disease genes were taken from the HuGE database of published GWAS genes (44), with a total of 2,711 diseases. HF related diseases were filtered out using keywords ‘heart’, ‘cardi’, ‘hypert’, ‘aort’, ‘fibro’.

### Pathways

Pathways were taken from MSigDB v3.1 (45) and Wikipathways (46), with a total of 8,690 sets of genes. A group of 106 genes corresponding to a previously published Cardiac Hypertrophic Signaling Network (CHSN) (20) was added under the name “ SAUCERMAN_cardiac_hypertrophy_pathway”.

### TF enrichment

The cytoscape plugin iRegulon (30) was used to predict putative upstream TF regulating the studied sets of genes. Default parameters were used: 9,713 PWMs scanning 20kb centered around TSS.

### Computation of statistics

All statistics (correlations, t-test, Wilcoxon test, hypergeometric test) were computed using R. Hierarchical clustering was performed using default parameters of the R hclust function. Z scores correspond to the number of standard deviations a given observation is away from the mean of the null (random) distribution and are computed as follow:

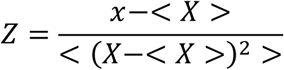
where x is the observed value, X is a set of random predictions, and <.> denotes the average.

### Cell Culture and Treatments

Right ventricular myocytes were isolated and cultured, as reported (47) using 2-4 day old rats. Myocytes and fibroblasts were separated with Percoll density gradient. For knockdown experiments cells were transfected with *Hes1* siRNA using lipofectamine RNAimax (life technologies).

### RNA Isolation and qPCR

RNA isolation from cells was performed using Qiazol lysis reagent. cDNA synthesis was performed using the High Capacity Reverse Transcription cDNA Kit (Life Technologies). qPCR was performed using the LightCycler 480 (Roche). The number of replicates per condition is shown in Supplementary table S3, with values ranging from 6 to 9.

### Quantification of cardiomyocyte cell size

Quantification of cardiomyocyte cell size was done following transfection with either control or Hes1 siRNA and a 48 hour treatment with control or isoproterenol or phenylepherine containing media. Images were taken on a Nikon Eclipse TE2000-U microscope. Images were analyzed using the Nikon Imagine System (NIS). 150 cells were used to compute the SEM.

## Acknowledgements

This research was supported by NIH/NHLBI grants 5R01HL114437-02 and 5R01HL05242 and by the Laubisch and Kawata Endowments.

---

**Figure S1.**
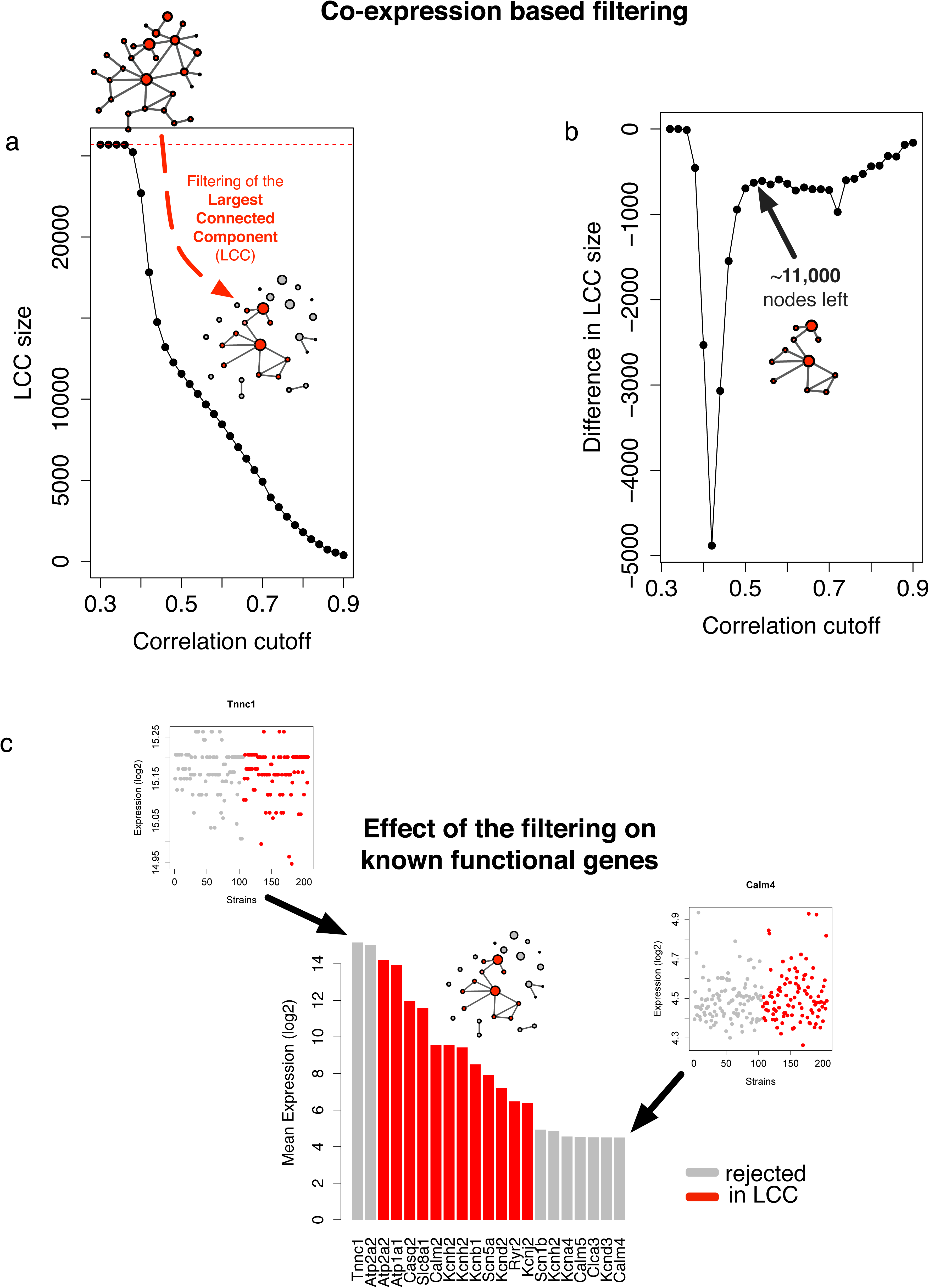
A network approach to expression filtering. a. A Pearson correlation co-expression network of the genes FCs is constructed. Links with absolute value smaller than a given cutoff are eliminated, and the Largest Connected Component (LCC) size is computed. A strong LCC size decrease is observed around a cutoff of 0.4.
b. The derivative of the previous plot is shown. The cutoff of 0.5 corresponding to the end of the drop-off is selected to define 11,279 genes in the LCC.
c. Examples of cardiac functionally relevant genes in and out of the LCC. Genes left out have either low expression (*Calm4*) or saturation issues (*Tnnc1*) which introduce noise in the correlation process.

**Figure S2.**
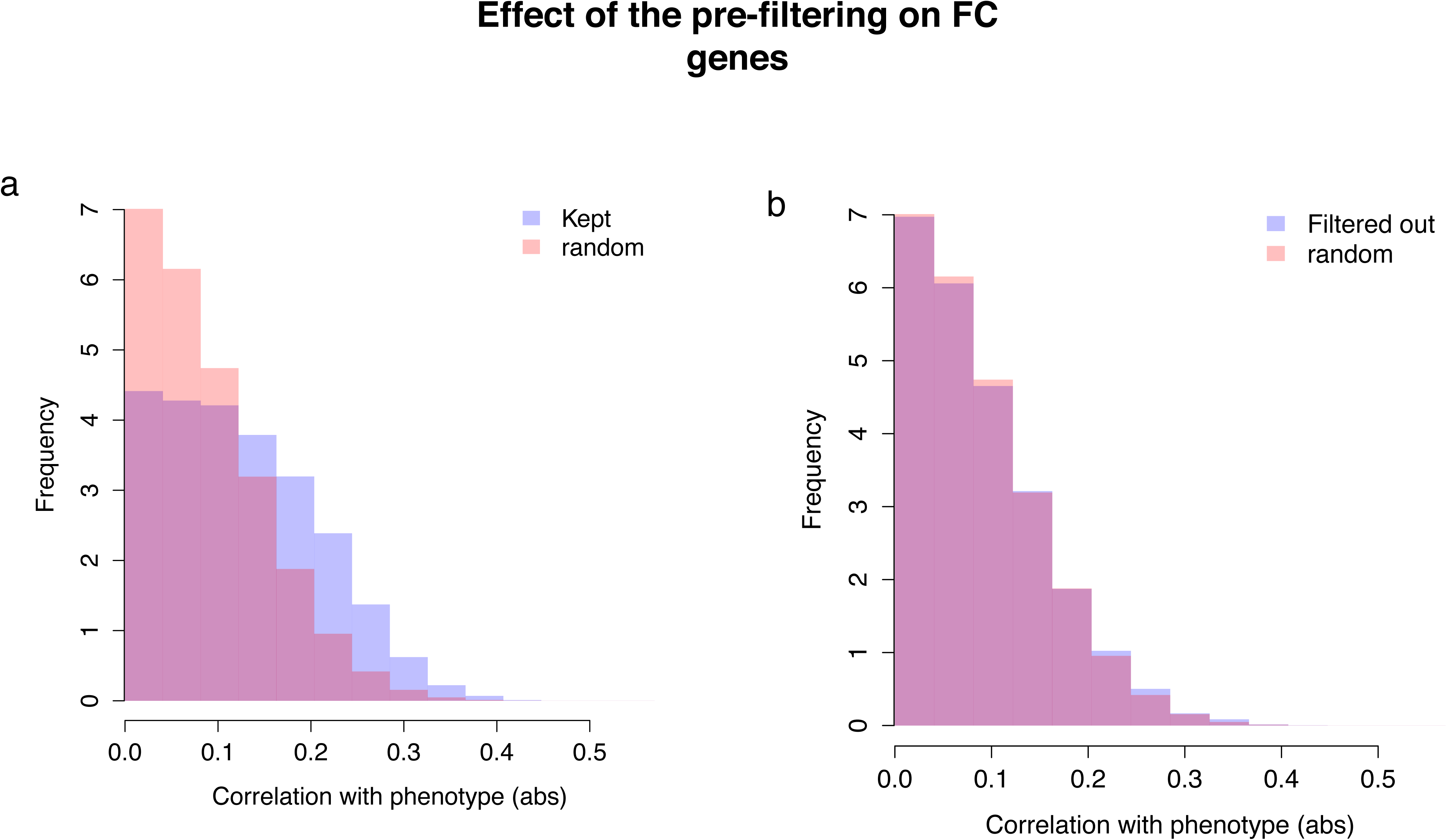
Effect of the pre-filtering on FC genes. Same plots than Figure 2a, using the filtered set of genes (a), or the genes filtered out by the method from Figure S1 (b). This shows that the genes highly correlated to hypertrophy are extracted during the filtering process, while genes with low correlations are filtered out.

**Figure S3.**
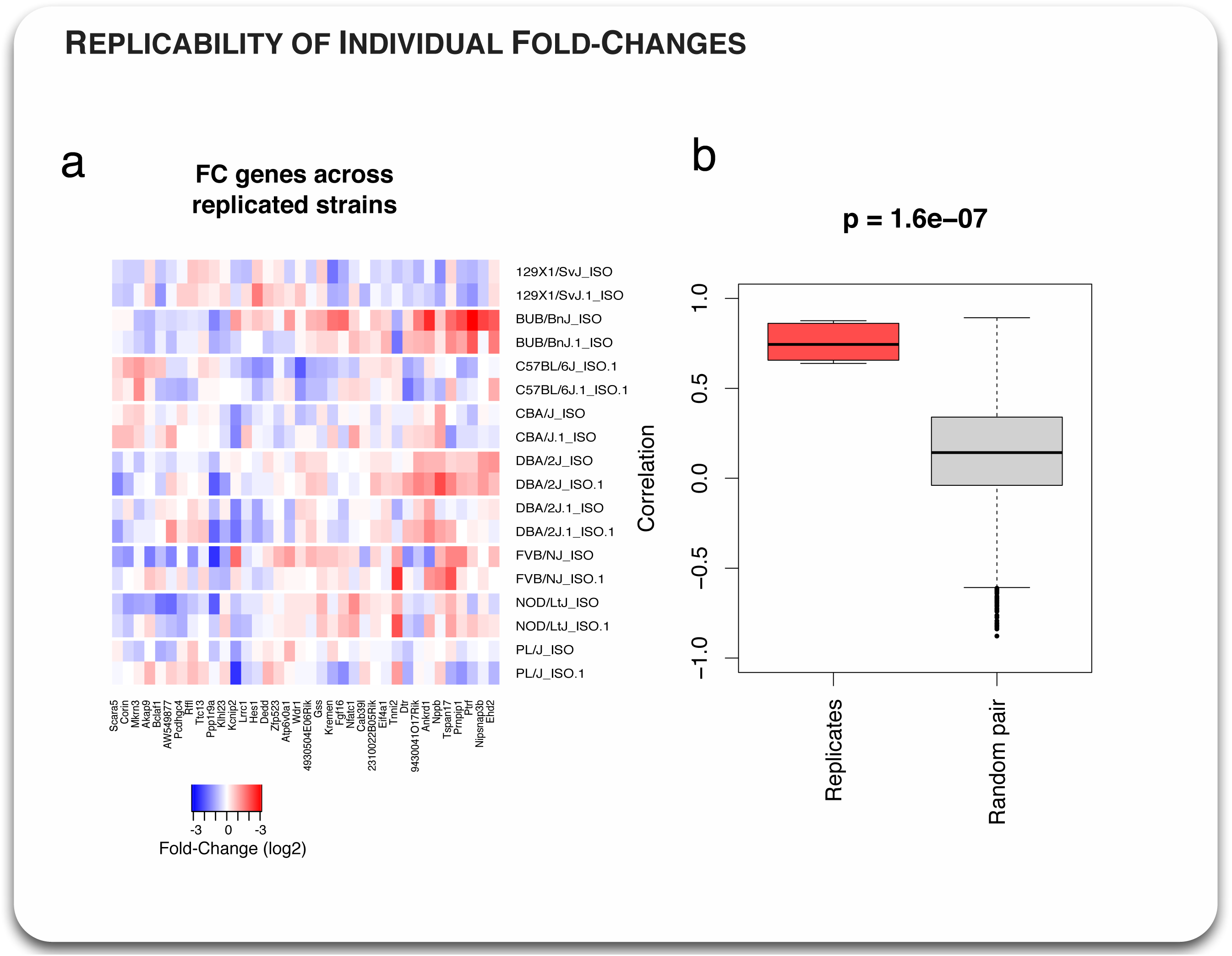
Replicability of individual fold-changes. a. Heatmap showing the fold-changes of the 36 hypertrophic genes for pairs of mice from 9 replicated strains.
b. The quality of replication is assessed by computing the Spearman correlation between the fold-changes of the 36 genes between replicated strains (red) and for random pairs of non-replicated strains (gray). The average correlation is 0.76, which is significantly higher than between two distinct strains where it is 0.14 (p=1.6e-7, Wilcoxon test).

**Figure S4.**
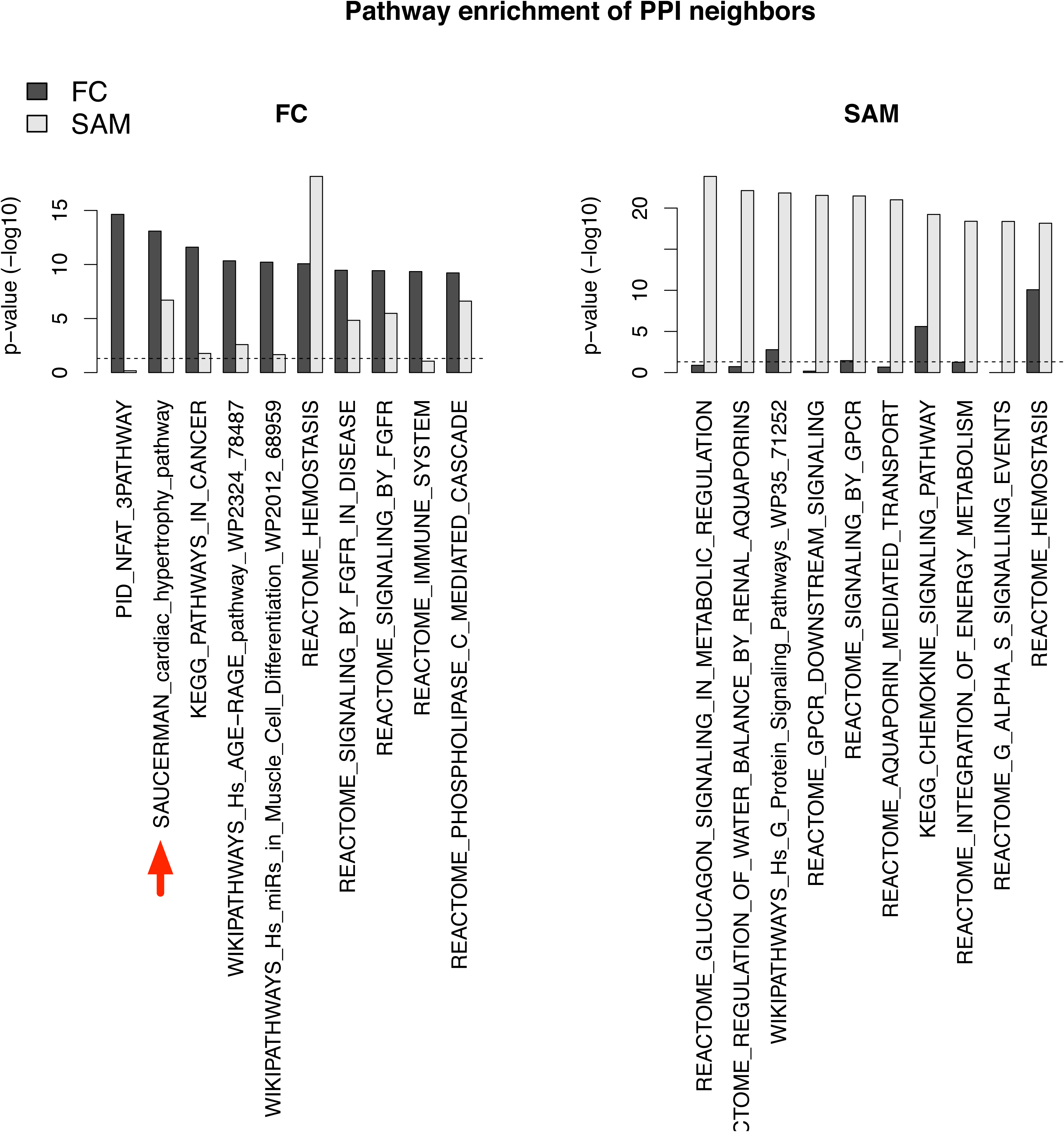
Pathway enrichment of neighbor genes in the interactome. a,b. Pathway enrichment was assessed on first neighbor genes of the FC (a) and SAM (b) genes in the human Protein-Protein Interactome (33). The 10 most enriched pathways are shown. FC neighbors are seen to be highly enriched in a previously published Cardiac Hypertrophy Signaling Network or CHSN (20).

**Figure S5.**
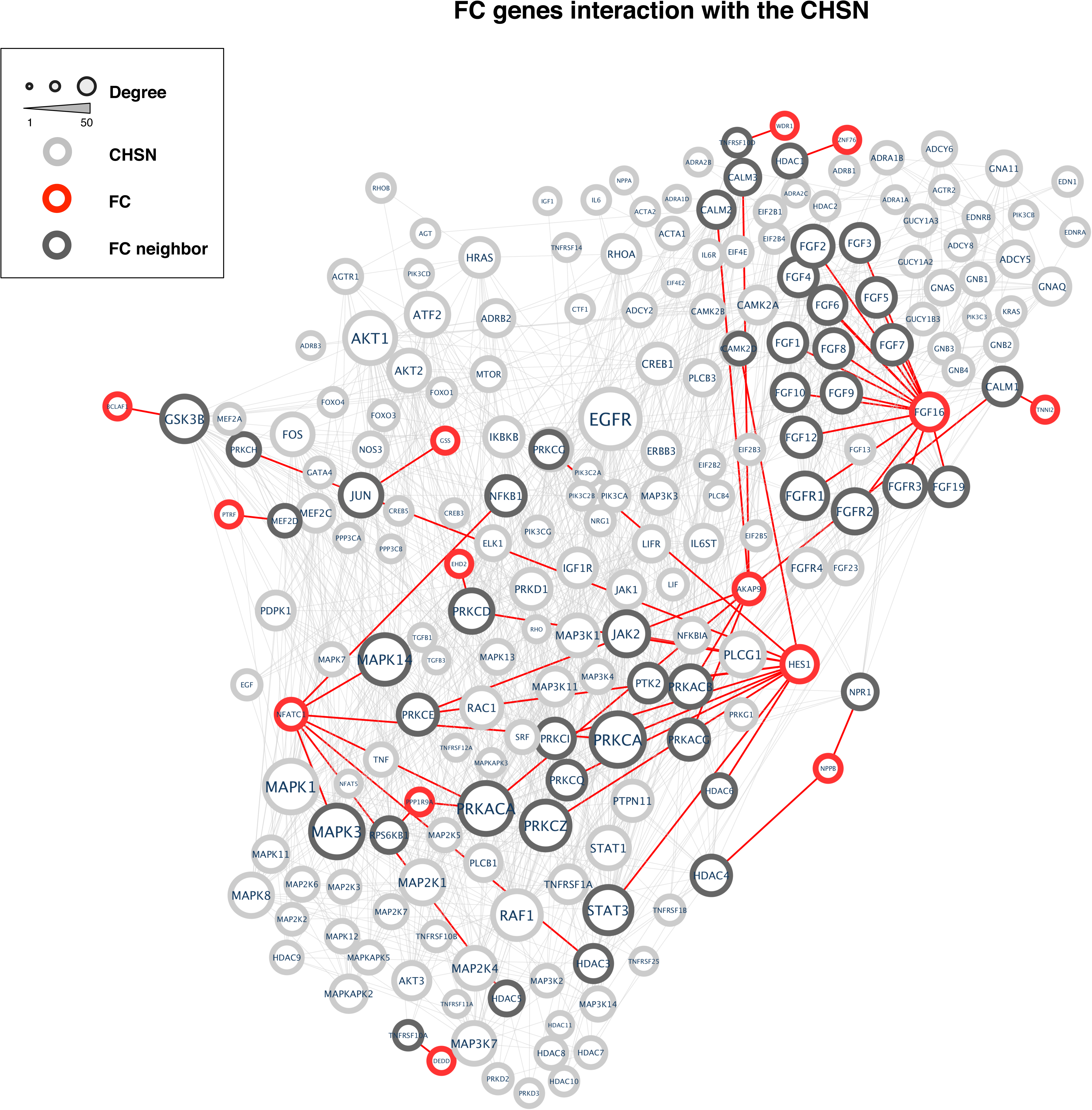
FC genes interaction with the CHSN. Network visualization of the CHSN (gray nodes) and their first neighbors in the FC set (red) in the interactome (33). Darker nodes from CHSN indicate interaction with a FC protein.

**Figure S6.**
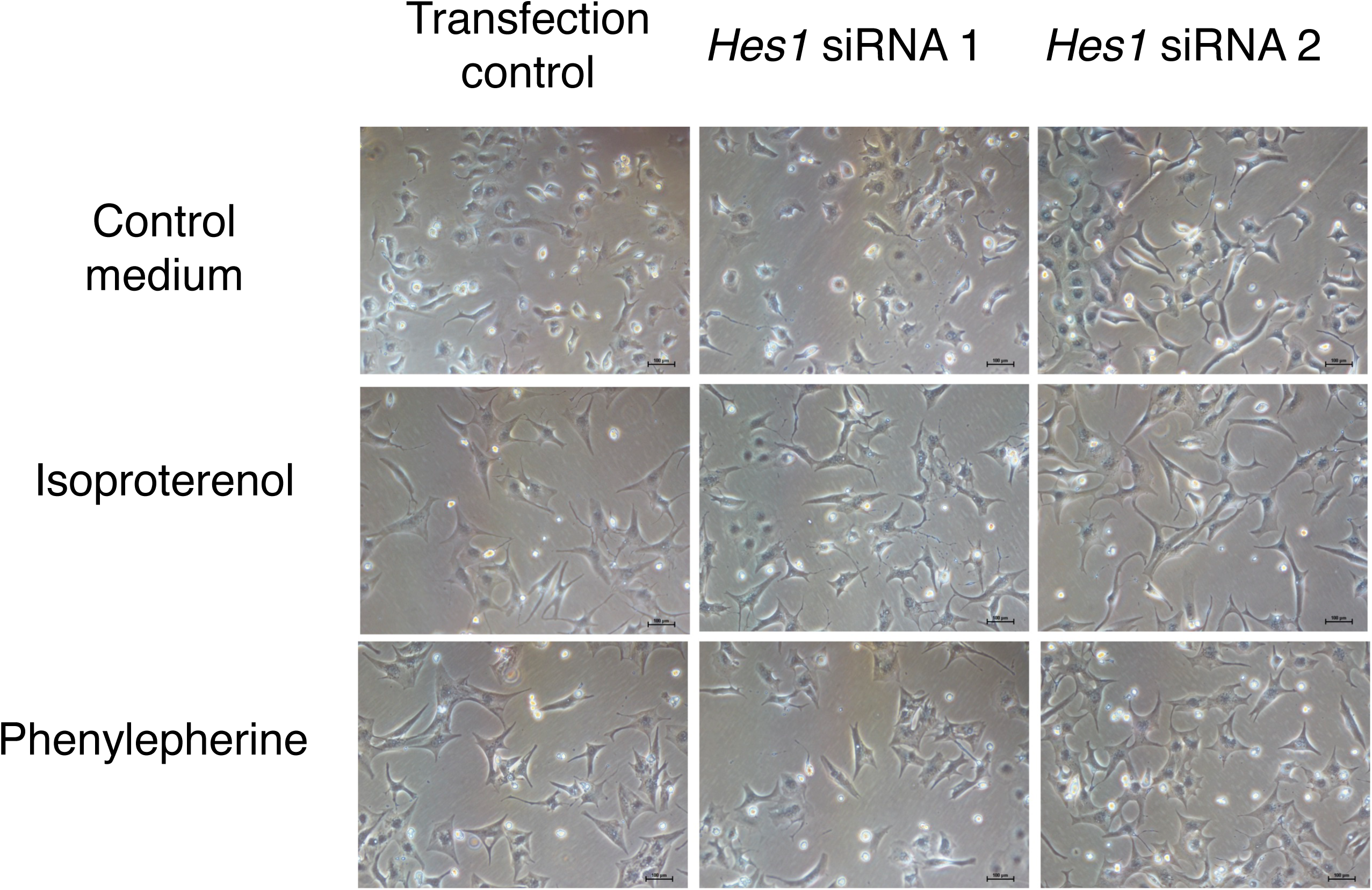
Microscopic images of *Hes1* siRNA assays in neonatal cardiac cell size. Microscopic images of neonatal rat cardiomyocytes following transfection with either control or Hes1 siRNAs and a 48 hour treatment with control or isoproterenol or phenylepherine containing media. Scale bars show 100 μm.

**Figure S7.**
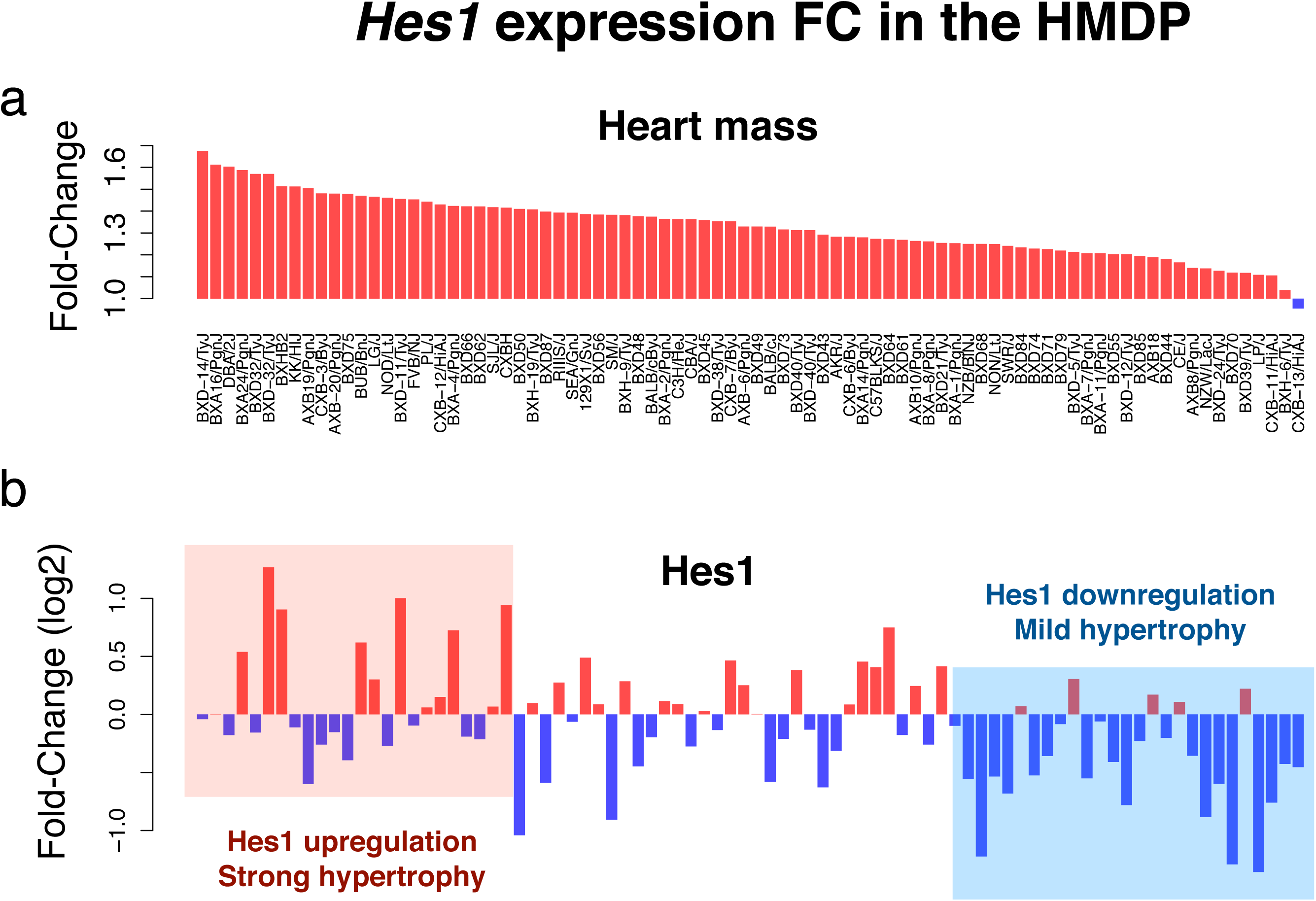
*Hes1* in the HMDP. Barplots comparing Heart mass (a) and *Hes1* (b) fold-changes across HMDP strains as in Figure 1h,k. Strains with no or mild hypertrophy show a negative fold-change of *Hes1*, an effect consistent with the *Hes1* siRNA assay in neonatal rat cells of Figure 4c.

## References

1. Albert FW & Kruglyak L (2015) The role of regulatory variation in complex traits and disease. Nature reviews. Genetics.

2. Bui AL, Horwich TB, & Fonarow GC (2011) Epidemiology and risk profile of heart failure. Nat Rev Cardiol 8(1): 30–41.

3. Cambronero F, et al. (2009) Biomarkers of pathophysiology in hypertrophic cardiomyopathy: implications for clinical management and prognosis. Eur Heart J 30(2): 139–151.

4. Heineke J & Molkentin JD (2006) Regulation of cardiac hypertrophy by intracellular signalling pathways. Nat Rev Mol Cell Biol 7(8): 589–600.

5. Blaxall BC, Spang R, Rockman HA, & Koch WJ (2003) Differential myocardial gene expression in the development and rescue of murine heart failure. Physiol Genomics 15(2): 105–114.

6. Gao Z, et al. (2008) Key pathways associated with heart failure development revealed by gene networks correlated with cardiac remodeling. Physiol Genomics 35(3): 222–230.

7. Asakura M & Kitakaze M (2009) Global gene expression profiling in the failing myocardium. Circ J 73(9): 1568–1576.

8. Weiss JN, et al. (2012) “Good enough solutions” and the genetics of complex diseases. Circ Res 111(4): 493–504.

9. Taylor AL, Hickey TJ, Prinz AA, & Marder E (2006) Structure and visualization of high-dimensional conductance spaces. J Neurophysiol 96(2): 891–905.

10. Salari K, Watkins H, & Ashley EA (2012) Personalized medicine: hope or hype? Eur Heart J 33(13): 1564–1570.

11. Creemers EE, Wilde AA, & Pinto YM (2011) Heart failure: advances through genomics. Nature reviews. Genetics 12(5): 357–362.

12. Ghazalpour A, et al. (2012) Hybrid mouse diversity panel: a panel of inbred mouse strains suitable for analysis of complex genetic traits. Mamm Genome 23(9-10):680–692.

13. Rau CD, et al. (2015) Mapping genetic contributions to cardiac pathology induced by Beta-adrenergic stimulation in mice. Circ Cardiovasc Genet 8(1): 40–49.

14. Lin H, et al. (2014) Gene expression and genetic variation in human atria. Heart Rhythm 11(2): 266–271.

15. van den Borne SW, et al. (2009) Mouse strain determines the outcome of wound healing after myocardial infarction. Cardiovasc Res 84(2): 273–282.

16. Shah AP, et al. (2010) Genetic background affects function and intracellular calcium regulation of mouse hearts. Cardiovasc Res 87(4): 683–693.

17. Barrick CJ, Rojas M, Schoonhoven R, Smyth SS, & Threadgill DW (2007) Cardiac response to pressure overload in 129S1/SvImJ and C57BL/6J mice: temporal- and background-dependent development of concentric left ventricular hypertrophy. Am J Physiol Heart Circ Physiol 292(5):H2119–2130.

18. Kiper C, Grimes B, Van Zant G, & Satin J (2013) Mouse strain determines cardiac growth potential. PLoS One 8(8):e70512.

19. Tusher VG, Tibshirani R, & Chu G (2001) Significance analysis of microarrays applied to the ionizing radiation response. Proceedings of the National Academy of Sciences of the United States of America 98(9): 5116–5121.

20. Ryall KA, et al. (2012) Network reconstruction and systems analysis of cardiac myocyte hypertrophy signaling. J Biol Chem 287(50): 42259–42268.

21. Callow MJ, Dudoit S, Gong EL, Speed TP, & Rubin EM (2000) Microarray expression profiling identifies genes with altered expression in HDL-deficient mice. Genome Res 10(12): 2022–2029.

22. Ritchie ME, et al. (2015) limma powers differential expression analyses for RNA-sequencing and microarray studies. Nucleic Acids Res 43(7):e47.

23. Jin H, et al. (2010) KChIP2 attenuates cardiac hypertrophy through regulation of Ito and intracellular calcium signaling. Journal of molecular and cellular cardiology 48(6): 1169–1179.

24. Kuo HC, et al. (2001) A defect in the Kv channel-interacting protein 2 (KChIP2) gene leads to a complete loss of I(to) and confers susceptibility to ventricular tachycardia. Cell 107(6): 801–813.

25. Grubb S, et al. (2014) Loss of K+ currents in heart failure is accentuated in KChIP2 deficient mice. J Cardiovasc Electrophysiol 25(8): 896–904.

26. Bignolais O, et al. (2011) Early ion-channel remodeling and arrhythmias precede hypertrophy in a mouse model of complete atrioventricular block. Journal of molecular and cellular cardiology 51(5): 713–721.

27. Fan D, Takawale A, Lee J, & Kassiri Z (2012) Cardiac fibroblasts, fibrosis and extracellular matrix remodeling in heart disease. Fibrogenesis Tissue Repair 5(1):15.

28. Baudino TA, Carver W, Giles W, & Borg TK (2006) Cardiac fibroblasts: friend or foe? Am J Physiol Heart Circ Physiol 291(3):H1015–1026.

29. Gardner DG (2003) Natriuretic peptides: markers or modulators of cardiac hypertrophy? Trends Endocrinol Metab 14(9): 411–416.

30. Janky R, et al. (2014) iRegulon: from a gene list to a gene regulatory network using large motif and track collections. PLoS Comput Biol 10(7):e1003731.

31. Paul V, et al. (2014) Scratch2 modulates neurogenesis and cell migration through antagonism of bHLH proteins in the developing neocortex. Cereb Cortex 24(3): 754–772.

32. Wu-Wong JR (2011) Vitamin D therapy in cardiac hypertrophy and heart failure. Curr Pharm Des 17(18): 1794–1807.

33. Menche J, et al. (2015) Disease networks. Uncovering disease-disease relationships through the incomplete interactome. Science 347(6224):1257601.

34. Molkentin JD (2004) Calcineurin-NFAT signaling regulates the cardiac hypertrophic response in coordination with the MAPKs. Cardiovasc Res 63(3): 467–475.

35. Benjamin IJ, et al. (1989) Isoproterenol-induced myocardial fibrosis in relation to myocyte necrosis. Circ Res 65(3): 657–670.

36. Ho CY, et al. (2010) Myocardial fibrosis as an early manifestation of hypertrophic cardiomyopathy. N Engl J Med 363(6): 552–563.

37. Tamura N, et al. (2000) Cardiac fibrosis in mice lacking brain natriuretic peptide. Proceedings of the National Academy of Sciences of the United States of America 97(8): 4239–4244.

38. Rau CD, et al. (2017) Systems Genetics Approach Identifies Gene Pathways and Adamts2 as Drivers of Isoproterenol-Induced Cardiac Hypertrophy and Cardiomyopathy in Mice. Cell Syst 4(1):121–128 e124.

39. de la Pompa JL (2009) Notch signaling in cardiac development and disease. Pediatr Cardiol 30(5): 643–650.

40. de la Pompa JL & Epstein JA (2012) Coordinating tissue interactions: Notch signaling in cardiac development and disease. Dev Cell 22(2): 244–254.

41. Zhou XL, Zhao Y, Fang YH, Xu QR, & Liu JC (2014) Hes1 is upregulated by ischemic postconditioning and contributes to cardioprotection. Cell Biochem Funct 32(8): 730–736.

42. Gao Z, et al. (2006) Transcriptomic profiling of the canine tachycardia-induced heart failure model: global comparison to human and murine heart failure. Journal of molecular and cellular cardiology 40(1): 76–86.

43. Ruiz P & Witt H (2006) Microarray analysis to evaluate different animal models for human heart failure. in Journal of molecular and cellular cardiology, pp 13–15.

44. Yu W, Gwinn M, Clyne M, Yesupriya A, & Khoury MJ (2008) A navigator for human genome epidemiology. Nature genetics 40(2): 124–125.

45. Liberzon A, et al. (2011) Molecular signatures database (MSigDB) 3.0. Bioinformatics 27(12): 1739–1740.

46. Kelder T, et al. (2012) WikiPathways: building research communities on biological pathways. Nucleic Acids Res 40(Database issue): D1301–1307.

47. Brown DA, et al. (2005) Modulation of gene expression in neonatal rat cardiomyocytes by surface modification of polylactide-co-glycolide substrates. J Biomed Mater Res A 74(3): 419–429.

48. Ryall KA, et al. (2012) Network Reconstruction and Systems Analysis of Cardiac Myocyte Hypertrophy Signaling. in Journal of Biological Chemistry, pp 42259–42268.

49. Gopalakrishnan K, et al. (2011) Augmented rififylin is a risk factor linked to aberrant cardiomyocyte function, short-QT interval and hypertension. Hypertension 57(4): 764–771.

50. Yuan B, et al. (2014) A cardiomyocyte-specific Wdr1 knockout demonstrates essential functional roles for actin disassembly during myocardial growth and maintenance in mice. The American journal of pathology 184(7): 1967–1980.

51. Wallen T, Landahl S, Hedner T, Nakao K, & Saito Y (1997) Brain natriuretic peptide predicts mortality in the elderly. Heart 77(3): 264–267.

52. Wei Z, et al. (2011) A common genetic variant in the 3′-UTR of vacuolar H+-ATPase ATP6V0A1 creates a micro-RNA motif to alter chromogranin A processing and hypertension risk. Circ Cardiovasc Genet 4(4): 381–389.

53. Bogomolovas J, et al. (2015) Induction of Ankrd1 in Dilated Cardiomyopathy Correlates with the Heart Failure Progression. Biomed Res Int 2015: 273936.

54. Iwamoto R, et al. (2003) Heparin-binding EGF-like growth factor and ErbB signaling is essential for heart function. Proceedings of the National Academy of Sciences of the United States of America 100(6): 3221–3226.

55. Rochais F, et al. (2009) Hes1 is expressed in the second heart field and is required for outflow tract development. PLoS One 4(7):e6267.

56. de Villiers CP, et al. (2014) AKAP9 is a genetic modifier of congenital long-QT syndrome type 1. Circ Cardiovasc Genet 7(5): 599–606.

57. Meune C, et al. (2012) Blood glutathione decrease in subjects carrying lamin A/C gene mutations is an early marker of cardiac involvement. Neuromuscul Disord 22(3): 252–257.

58. Damy T, et al. (2009) Glutathione deficiency in cardiac patients is related to the functional status and structural cardiac abnormalities. PLoS One 4(3):e4871.

59. Adamy C, et al. (2005) Tumor necrosis factor alpha and glutathione interplay in chronic heart failure. Arch Mal Coeur Vaiss 98(9): 906–912.

60. Zhao YY, et al. (2002) Defects in caveolin-1 cause dilated cardiomyopathy and pulmonary hypertension in knockout mice. Proceedings of the National Academy of Sciences of the United States of America 99(17): 11375–11380.

61. Laurell T, et al. (2014) Identification of three novel FGF16 mutations in X-linked recessive fusion of the fourth and fifth metacarpals and possible correlation with heart disease. Mol Genet Genomic Med 2(5): 402–411.

62. Gudmundsson H, et al. (2010) EH domain proteins regulate cardiac membrane protein targeting. Circ Res 107(1): 84–95.

63. Lopes LR, et al. (2013) Genetic complexity in hypertrophic cardiomyopathy revealed by high-throughput sequencing. J Med Genet 50(4): 228–239.

64. Cordero P, et al. (2016) A community overlap strategy reveals central genes and networks in heart failure. bioRxiv.

65. Nakamura T, Nakamura T, & Matsumoto K (2008) The functions and possible significance of Kremen as the gatekeeper of Wnt signalling in development and pathology. J Cell Mol Med 12(2): 391–408.

66. van de Schans VA, et al. (2007) Interruption of Wnt signaling attenuates the onset of pressure overload-induced cardiac hypertrophy. Hypertension 49(3): 473–480.

67. Wang W, et al. (2012) Salt-sensitive hypertension and cardiac hypertrophy in transgenic mice expressing a corin variant identified in blacks. Hypertension 60(5):1352–1358.

